# Region-specific associations between cerebral arterial elasticity and grey matter volume: Evidence for lateral prefrontal vulnerability

**DOI:** 10.64898/2025.12.03.691980

**Authors:** Nicholas Ware, Jenna Johnson, Monica Fabiani, Sarah Johnson, Patricia Michie, Kathy Low, Christian Behler, Bryan Paton, Daniel Barker, Gabriele Gratton, Ashleigh E. Smith, Frini Karayanidis

**Author notes:** **Corresponding authors:** Nicholas Ware, Frini Karayanidis, **Address:** School of Psychological Sciences, College of Engineering, Science and Environment, University of Newcastle, University Dr, Callaghan NSW 2308, Australia.

## Abstract

**Background:** Ageing is associated with increased cardiovascular health risks and disproportionate atrophy in frontotemporal brain. Regional cerebral arterial elasticity correlates with regional grey matter volume, with stronger associations frontally and in older adults. Cardiorespiratory fitness (CRF) is linked to preserved brain structure and greater arterial elasticity, while sex differences exist in the timing of vascular versus structural changes. This study examines whether regional cerebral arterial elasticity in frontotemporal cortex mediates the association between age and corresponding grey matter volume decline, and whether sex and/or CRF moderate this relationship.

**Methods:** We analysed data from 162 healthy adults (60-70 years) with structural MRI and diffuse optical tomography of the cerebral arterial pulse (Pulse-DOT) from the ACTIVate cohort. CRF was estimated from demographic and physiological measures. Pulse Relaxation Function (PReFx), an optical index of regional arterial elasticity, was measured across 28 frontal, temporal, and parietal regions of interest (ROI). Grey matter (GM) volume was quantified for corresponding ROIs. Mediation and moderated mediation models tested whether PReFx mediated the relationship between age and GM volume at bilateral frontal and temporal ROIs, and whether biological sex or CRF moderated this relationship.

**Results:** Regional bivariate correlations identified associations between age, PReFx and GM volume across multiple ROIs, which PReFx and GM volume being associated primarily in left frontal and temporal areas. PReFx partially mediated the effect of age on GM volume in a left mid-inferior frontal ROI, accounting for approximately 16% of the total age effect and this relationship was only evident in females. Although higher CRF was associated with greater PReFx, it did not moderate the relationship between age, PReFx and GM volume.

**Conclusions:** Consistent with cascade models of neurovascular aging, cerebral arterial stiffening was found to partially explain the effect of increasing age on frontal GM volume, even in this highly age-restricted and high functioning cohort. This effect was only significant over the left prefrontal cortex, consistent with greater vulnerability of frontal brain and associated cognitive functions. It was also exclusively present in females, who had better cardiovascular health, larger grey matter volume and greater arterial elasticity than males. These findings are consistent with pulse-DOT measures of cerebral arterial elasticity being more sensitive to subclinical brain structural variability.

## INTRODUCTION

Aging is associated with subtle and progressive cortical atrophy, even in the absence of neurodegenerative disease (Cox & Deary, 2022). However, atrophy is not uniform across the cortex but varies across regions and individuals, with disproportionate atrophy in frontotemporal cortices (Fjell et al., 2013; Raz et al., 2010; Zanto & Gazzaley, 2019). This heterogeneity may be due, in part, to variability in emerging cardiovascular deterioration, especially regional changes in cerebral arterial elasticity (Chiarelli et al., 2017; Fabiani et al., 2014; Tan et al., 2019). Cardiorespiratory fitness (CRF) is a modifiable lifestyle factor that is associated with cardiovascular health and with peripheral and whole-brain measures of arterial elasticity (Boreham et al., 2004; Fabiani et al., 2014; Jae et al., 2024). However, the extent to which CRF explains variability in regional cerebral elasticity, and whether it influences the relationship between cerebral arterial elasticity and cortical atrophy, remain unclear.

The walls of elastic arteries thicken and stiffen with increasing age (Mitchell et al., 2011). This arterial stiffness limits the buffering capacity of the arterial system and increases both the velocity and pulsatile component of blood flow. Arterial stiffening is associated with the presence and severity of age-related cardiovascular disease risk factors (e.g., hypertension, diabetes mellitus; AlGhatrif et al., 2013; Zheng et al., 2020) and is an important predictor of several clinical cardiovascular health conditions (e.g., stroke, myocardial infarction and cerebral small vessel disease; see Singer et al., 2014 for review).

High-flow, low-resistance organs, such as the brain, are particularly vulnerable to arterial stiffening (Stone et al., 2015). Under normal physiological conditions, large cerebral arteries (e.g., middle and anterior cerebral arteries) buffer pulsatile flow before it reaches the microvasculature. Within the cranial cavity, elastic arterial walls dampen pulsatile flow (Windkessel effect; Westerhof et al., 2009), while increases in arterial blood volume are offset by displacement of CSF and venous blood from the rigid intracranial space (Monro-Kellie doctrine; Monro, 1783).As arterial elasticity declines, pulsatile energy is transmitted more forcefully into the vascular bed, damaging the microcirculation that supplies brain parenchyma. This initiates a cascade of downstream processes, including blood–brain barrier disruption, impaired cerebrovascular reactivity, and regional hypoperfusion. Microvascular injury is, in turn, associated with parenchymal deterioration, manifesting as grey matter atrophy, white matter hyperintensities and axonal degeneration (see Zimmerman et al., 2021 for review).

There is strong evidence that arterial elasticity is associated with cerebral structural damage, with most evidence to date coming from systemic measures of arterial elasticity (e.g., pulse wave velocity (PWV), pulse pressure) and central measures derived from transcranial Doppler (Badji et al., 2019; Singer et al., 2014, p. 20). For instance, a recent study by Won et al., (2025) showed extensive relationships between age, carotid-femoral PWV, and measures of brain structural health, including white matter hyperintensities, microstructural organization, and grey matter volume, both globally and regionally. However, these relationships are not uniform across the brain, e.g., areas at the edge of arterial watersheds are particularly susceptible to effects of hypoxia and ischemia (Kapasi et al., 2018). Frontotemporal cortices are disproportionately vulnerable to vascular-related degeneration (Raz et al., 2007), and several studies have shown regional specificity in these arterial-structural associations. Yu et al., (2025) demonstrated that in older adults, associations between carotid-femoral PWV and frontal lobe structure were strongest in left lateral hemisphere prefrontal cortices. Similarly, Pasha et al., (2015) reported that, in middle-aged adults, systemic arterial stiffness was associated with reduced cortical thickness in left IFG.

The rate of age-related decline in arterial elasticity is strongly and inversely correlated with CRF, a systemwide measure of cardiovascular efficiency (Ross et al., 2016). Higher CRF is associated with greater arterial elasticity in both cardiac and cerebral vasculature (Albin et al., 2020; Clements et al., 2024; Jae et al., 2024), lower prevalence of cardiovascular disease risk factors (Sloan, 2024), and enhanced cerebral perfusion (Zimmerman et al., 2014). High CRF is also associated with lower frontotemporal cortical atrophy (Erickson et al., 2022). This relationship is mediated, in part, by the upregulation of neurotrophic and vascular growth factors that promote angiogenesis and support neurovascular health (Davenport et al., 2012).

However, much of the evidence linking CRF and arterial stiffness with brain structure has relied on measures taken outside of the skull (e.g., from aortic or carotid-femoral pulse wave velocity assessments; Qiu et al., 2003; Waldstein et al., 2008). These measures infer the status of cerebral arteries from that of extracranial arteries, precluding direct characterization of intracranial arterial health. While there are methods to assess the health of arteries more proximal to or within the brain, they cannot quantify regional variation in arterial elasticity. For instance, carotid sonography evaluates arterial wall structure and plaque presence, but offers limited insight into intracranial arteries (Silvestrini et al., 2011). Transcranial Doppler (TCD) sonography measures intracranial arterial elasticity (e.g., Pulsatility or Resistance Index, (Markus, 2000) from only a few specific insonation points viewed only through acoustic windows. In addition, TDC cannot be performed on all older individuals due to calcification of the temporal bone, which leads to transtemporal window failure (Chan et al., 2023; He et al., 2022).

Fabiani, Gratton and Low developed a measure of regional cortical arterial elasticity by analysing the shape and timing of the cardiac component of the intracranial near-infrared waveform (i.e., diffuse optical tomography of the arterial pulse, Pulse-DOT) across different spatial locations of the cortical mantle (Fabiani et al., 2014). The *pulse relaxation function* (PReFx; formerly labelled *arterial compliance* in Fabiani et al., 2014) measures the shape of the arterial pulse waveform between the forward moving systolic pressure wave and the reflected pressure wave. Variation in the temporal separation of these waves provides information about the *elasticity of cerebral arteries.* Higher values of PReFx indicate a greater temporal separation, indicative of lower cerebral arterial stiffness.

Global PReFx (i.e., mean value across the cortex) is inversely associated with age and positively associated with both cardiorespiratory fitness and total brain, grey and white matter volumes (Fabiani et al., 2014; Tan et al., 2017). When measured regionally (i.e., mean PReFx across all voxels within an individual ROI), Chiarelli et al. (2017) reported significant spatial correlations between regional cerebral arterial elasticity and cortical volume. Whole-brain analysis across 50 Desikan–Killiany atlas ROIs in a small lifespan sample (n = 48, aged 18-75 years) showed that regions with lower PReFx, indicating reduced arterial elasticity, also had lower cortical volume. PReFx – cortical volume associations were strongest in frontotemporal cortical areas and only among older adults (> 48 years; see Chiarelli et al., 2017 Figure 3). These findings suggest that regional measures of arterial elasticity may offer greater explanatory power than global indices and that cerebral arterial stiffening may contribute to spatial variability in cortical atrophy. Importantly, they also indicate that cerebral arterial elasticity does not decline uniformly across the cortex but differentially in vulnerable cortical regions. Specifically, subtle and progressive reduction in frontotemporal arterial elasticity may contribute to the differential vulnerability of these regions to age-related atrophy, hypertensive injury, cerebral small vessel disease, and vascular cognitive impairment (Kalaria et al., 2024; Kuller et al., 2009; Raz et al., 2007). However, Chiarelli et al.’s older adult subgroup that showed these associations between arterial elasticity and measures of cortical structure was relatively small (n = 24) and had a broad age range (48-75 years). This may have obscured more specific relationships within older adults who are likely to show stronger vascular-structural coupling. Using a similar optical measure of regional cerebral arterial elasticity (i.e., pulse-transit-to-the-peak-of-the-pulse), Mohammadi et al., (2021) reported associations between arterial elasticity and cortical thickness in older adults (n = 30, aged 65-75 years) localised bilaterally to frontotemporal regions.

Bowie et al. (2024) showed sex differences in the trajectory of arterial changes. Specifically, in women, reduced arterial elasticity emerged around 55 years of age, and approximately 8-10 years earlier than structural brain changes. However, the factors that moderate this relationship between age, sex, arterial elasticity and brain structure remain unclear. It is possible that lifestyle factors that impact cardiorespiratory fitness may protect against age-related arterial stiffening (Erickson et al., 2014; Myers et al., 2015).

This study characterises the relationship between regional arterial elasticity (PReFx) and grey matter volume in a large cohort of cognitively healthy older adults with an age-restricted range (60-70 years) to minimize variability related to ageing. Arterial elasticity and grey matter volume were measured over frontal and temporal cortices, which are particularly vulnerable to ageing and cardiovascular health factors. Specifically, we first examined whether regional arterial elasticity mediates the relationship between age and grey matter volume, to replicate the above findings in a large age-restricted sample. We then examined whether cardiorespiratory fitness (CRF) moderates the relationship between age and arterial elasticity, buffering against age-related stiffening, and whether sex moderates the relationship between arterial elasticity and grey matter volume, given prior evidence for sex-specific stiffening trajectories (Bowie et al., 2024).

## METHODS

### Ethics

Phase 1 (baseline) data from the Newcastle cohort of the longitudinal ACTIVate Study (Smith et al., 2022) were included because optical imaging data were available. Data were collected between August 2020 and February 2022 at the University of Newcastle. The ACTIVate Study was registered with the Australia New Zealand Clinical Trials Registry (ACTRN12619001659190) on November 27, 2019. Ethical approval was obtained from the University of South Australia (202639) and subsequently registered with the University of Newcastle Human Research Ethics Committee (H-2019-0421).

### Participants

The cohort included 200 participants who met inclusion criteria for the ACTIVate Study (60–70 years, fluent in English). Participants were excluded if they reported a clinical diagnosis of a major neurological or psychiatric disorder, intellectual disability, major physical disability, mild cognitive impairment or dementia, or if they scored below 13/22 on the Montreal Cognitive Assessment (Blind; MoCA-B; Julayanont & Nasreddine, 2017) administered via telephone. The sample was predominantly White. The final sample included 162 participants, as 22 did not complete MRI and/or optical imaging, and another 16 had poor MRI or optical data quality.

### Measures

#### Socio-Demographic and Clinical Measures

Demographic and clinical information were gathered via a brief interview. Socioeconomic status (SES) was determined using the Index of Relative Socio-economic Advantage and Disadvantage (IRSAD), which draws on data from the Australian Census (ABS, 2023).

Physiological and anthropometric measures were collected following standard procedures (Smith et al., 2022). Systolic and diastolic blood pressure and resting heart rate were recorded as the average of three measures taken at five-minute intervals using a brachial electronic sphygmomanometer. Height and weight were measured for body mass index (BMI) calculation.

Venous blood samples were drawn following an overnight fast, centrifuged at 4000 rpm for 10 minutes, and plasma was aliquoted and frozen at −80°C until analysis. Prior to assay, samples were thawed on ice, vortexed, and centrifuged at 10,000 rpm for 2 minutes. Plasma glucose, total cholesterol, triglycerides, and HDL cholesterol were quantified using a KONELAB 20XTi automated analyzer with manufacturer-specific reagents (Thermo Fisher). LDL cholesterol was derived using the Friedewald equation: LDL = (total cholesterol - HDL) - (triglycerides/5).

#### Structural Magnetic Resonance Imaging (MRI)

A high-resolution structural MRI scan was acquired on a 3 Tesla Siemens PRISMA scanner using a 64-channel head and neck coil. T1-weighted brain images were collected via a 3D Magnetization Prepared Rapid Gradient Echo (MPRAGE) sequence (TR = 2,300 ms; TI = 900 ms; TE = 2.91 ms; flip angle = 9°; matrix size = 256 × 256 × 208; voxel size = 1 × 1 × 1 mm³; acquisition time = 5:12 min).

### Diffuse Optical Tomography of the Cerebral Arterial Pulse (Pulse-DOT)

#### Digitization and Co-registration

A Polhemus 3Space FASTRAK system (Polhemus, Colchester, VT) digitized all source and detector locations on the helmet, along with fiducial landmarks at the nasion and preauricular points. Three head-mounted receivers tracked minor head movements. Digitized coordinates were coregistered to the participant’s T1-weighted MRI using in-house software (Optimized Co-registration Package, OCP; Chiarelli et al., 2015). Participants were instructed to remain still during digitization and optical recording.

#### Optical Recording

Optical data were collected using a multi-channel frequency-domain oximeter (Imagent; ISS Inc., Champaign, IL, USA), featuring 16 time-multiplexed laser diodes and 4 detectors, yielding 128 source–detector (S-D) pairs per montage. Three helmet montages targeted the left, middle, and right prefrontal cortex in fixed order, totaling 384 S-D pairs. A soft foam helmet secured the optodes to the scalp. Each source delivered light at 830 and 690 nm (max 10 mW; ∼1 mW post-multiplex). Detector fibers transmitted received light to photomultiplier tubes (PMTs), modulated at 110.003125 MHz to produce a 3.125 kHz cross-correlation frequency. A Fast Fourier Transform of PMT output yielded DC intensity, AC intensity, and relative phase at 39.0625 Hz (25.6 ms resolution). Electrocardiogram (ECG) data were recorded at 1000 Hz (ADInstruments PowerLab Series 25) using electrodes placed on the left and right wrists.

#### Measurement of Pulse Parameters

Each montage (left, middle, right) was recorded in four 3-minute blocks (Rest 1, Breath-Holding 1, Breath-Holding 2, Rest 2). Breath-holding was used to measure changes in pulse amplitude as a measure of cardiovascular reactivity and are not analyzed here (Tan et al., 2016). We focused on AC intensity at 830 nm (Fabiani et al., 2014), yielding 64 channels per montage and 192 channels total. Source–detector distances (15–95 mm) allowed sampling of cortical depths up to ∼3 cm. Optical signals were scaled to their mean, band-pass filtered (0–10 Hz) and processed in Opt-3d (Gratton, 2000). Channels with 20-60 mm S-D distances were combined into voxel-wise waveforms, weighted by diffusion path length. Epochs were time-locked to the ECG R-wave, ensuring consistent sampling of the cardiac cycle across all locations.

### Data Processing

#### ECG Analysis

R-peaks were identified using a custom algorithm (1–50 Hz band-pass) in MATLAB R2021b (MathWorks), with visual inspection used to verify and correct misclassifications (e.g., large T waves). Optical pulse data were time-locked to the R-wave to ensure alignment across all channels.

#### Pulse Relaxation Function (PReFx)

PReFx, derived from each ROI using Opt-3d, quantifies pulse waveform shape between peak systole and diastole by subtracting a linear (triangular) component (Johnson et al., in preparation, Fig. 2). PReFx is normalised by the pulse amplitude (systolic peak = 1, diastolic peak = 0) and divided by the systole–diastole interval, making it independent of pulse amplitude and heart rate. A value of 0.5, which would reflect a constant decrease from the peak systole to the peak diastole, was subtracted from the value obtained above. Thus, positive values of PReFx reflect an accelerating descent from the peak systole (starting weak and becoming progressively stronger), whereas negative values reflect a decelerating descent (starting strong and becoming progressively weaker). In general, the more elastic the arteries, the higher the value of PReFx.

28 ROIs were defined a priori using AAL3 (Rolls et al., 2020), targeting frontotemporal and adjacent cortical regions (ACA and MCA supply). ROIs included frontal (precentral, middle/inferior frontal gyri), temporal (superior/middle/inferior temporal gyri, temporal poles), parietal (postcentral, supramarginal gyri), and midline regions (superior frontal gyri), restricted to superficial cortex (<3 cm depth) accessible to pulse-DOT (for ROI list, see Table 3).

Within Opt-3d, trials were excluded if: (1) systolic or diastolic peaks were undetected, (2) PReFx exceeded 1.0 or was below −0.2, (3) peak amplitude was <0.001 or >3 dB, (4) the interbeat interval (IBI) deviated from the ECG-derived IBI by >76.8 ms, or (5) IBI was <500 ms or >1500 ms. For each ROI, at least 10% of voxels were required to have ≥10 valid trials; ROIs not meeting this threshold were excluded. In total, 60 individual ROI-block combinations were removed across 26 participants (0.62% of the total dataset: 162 participants × 28 ROIs × 2 blocks). PReFx values were averaged across resting blocks 1 and 2 as they were highly correlated. Reliability for all pulse parameters is reported in Johnson et al. (in preparation).

#### Structural MRI

T1-weighted structural MRI scans were processed using Computational Anatomy Toolbox 12 (v12.9) for SPM12 (MATLAB R2023b) using the default volume workflow. Briefly, processing included tissue segmentation via Adaptive Maximum A Posteriori (AMAP) and Partial Volume Estimation (PVE) and spatial normalization using DARTEL aligned to the MNI152NLin2009cAsym template(Gaser et al., 2024). Regions of interest (ROIs) were defined *a priori* based on the AAL3 Atlas and focused on 28 frontal and frontal-adjacent ROIs (Table 2), matching those used for PReFx measures. All processed images underwent visual checks and CAT12’s automated quality metrics. All volumes were normalized against estimated total intracranial volume (eTIV) and expressed as a percentage for subsequent analyses (O’Brien et al., 2006).

#### Estimated Cardiorespiratory Fitness

Cardiorespiratory fitness was estimated using the non-exercise VO₂max prediction equation from (Jurca et al., 2005), which incorporates age, sex, BMI, and self-reported physical activity level. This approach avoids the need for maximal exercise testing and has predictive accuracy comparable to direct physiological testing (Ross & Myers, 2023).

Following Fabiani et al. (2014), Tan et al. (2019), and Clements et al. (2024), we removed the sex coefficient from the equation prior to analysis. This coefficient corrects for systematic sex differences in body size and lung capacity but does not represent meaningful individual variability in cardiovascular health.

### Statistical Analyses

#### Correlation Analyses

Zero-order Pearson correlations with 95% confidence intervals were estimated using 10,000 bootstrap iterations to assess: (1) age and regional arterial health (PReFx), and (2) age and grey matter (GM) volume across 28 ROIs, as well as (3) PReFx versus GM volume within corresponding ROIs.

#### Mediation Analyses

A simple mediation model (PROCESS Model 4) was fitted to establish whether regional arterial elasticity (PReFx) mediates the relationship between age and grey matter volume at bilateral frontal and temporal ROIs where regional arterial elasticity has been shown to be associated with measures of brain structure (Chiarelli et al., 2017, Mohammadi et al., 2021). A frontal and a temporal ROI were defined by averaging over three adjacent smaller ROI areas perfused by the left or right middle cerebral arteries (see Results for ROI details).

Where there was a significant mediation, moderated mediation models were fitted to identify factors that may moderate this effect. First, we examined whether cardiorespiratory fitness (CRF) moderates the relationship between age and both PReFx and GM volume (PROCESS Model 8). CRF has been shown to protect against both age-related arterial stiffening (Bowie et al., 2021; Clements et al., 2024; Zimmerman et al., 2021) and age-related GM volume decline (Erickson et al., 2014; Stillman et al., 2020; Voss et al., 2013) suggesting it may moderate both the age → PReFx pathway and the direct age → GM volume pathway. Second, we tested whether biological sex moderates the PReFx and GM volume relationship (PROCESS Model 14), as Bowie et al. (2024) reported sex-specific trajectories of arterial elasticity changes. In women, reduced arterial elasticity emerged earlier than structural brain changes.

Models were fitted in R (v4.3.1, RStudio v2024.09.0) using the PROCESS macro for R v5.0 (Hayes, 2022), with 10,000 bootstrap samples to 95% confidence intervals for indirect effects. The total variance accounted for by the mediator was quantified as a percentage of total variance [(indirect effect / total effect) × 100], where total effect is the sum of the direct effect and indirect effects. Visualizations used an MNI brain template rendered in surface (Rorden, 2025). Two-tailed p-values are reported unless otherwise noted.

Missing values included partially missing ROI-level PReFx values and a small number of incomplete demographic and physiological measures, were imputed using the missForest algorithm (see https://osf.io/s4vkw). This machine learning method predicts missing numerical and categorical values using random forest models until convergence (Stekhoven, 2025).

## RESULTS

### Cohort Characteristics

The cohort comprised 43% males, who were slightly older than the females (Table 1). On average, participants were highly educated, scored more than 1 SD above the mean on the NIH Crystallized Ability composite and had global cognitive functioning well above established cut-offs for mild cognitive impairment and dementia (88 and 82, respectively; Hsieh et al., 2013; Velayudhan et al., 2014). Average BMI was below the obesity cut-off (30 kg/m²).

**Table 1.** Demographic, cognitive and physiological characteristics of cohort, with sex comparisons.

| VARIABLES | OVERALL<br>n=162 | MALES<br>n=70 (43%) | FEMALES<br>n=92 (57%) | t-test<br>(2-tailed) |
| --- | --- | --- | --- | --- |
| Age | 65.4 (3.1) | 66.0 (3.3) | 64.9 (2.9) | **.038** |
| Education (years) | 16.9 (3.3) | 16.8 (3.3) | 16.9 (3.2) | .912 |
| NIH Crystallized Ability | 120 (11) | 119 (12) | 120 (10) | .685 |
| ACE III | 94 (4) | 94 (4) | 95 (3) | .122 |
| Est. Adj. Cardiorespiratory Fitness (METs) | 6.8 (1.9) | 6.9 (1.7) | 6.8 (2.0) | .883 |
| Body Mass Index (kg/m<sup>2</sup>) | 27.3 (5.1) | 27.8 (4.4) | 26.9 (5.5) | .257 |
| Systolic Blood Pressure (mmHg) | 140 (17) | 146 (16) | 137 (16) | **.001** |
| Diastolic Blood Pressure (mmHg) | 77 (8) | 79 (9) | 76 (8) | **.015** |
| Pulse Pressure (mmHg) | 63 (13) | 66 (12) | 61 (13) | **.010** |
| Resting Heart Rate (bpm) | 65 (11) | 63 (10) | 67 (11) | **.018** |
| Glucose (mg/dL) | 87.3 (10.8) | 90.3 (10.5) | 85.0 (10.5) | **< .01** |
| HDL Cholesterol (mg/dL) | 66.0 (19.2) | 59.8 (17.5) | 70.8 (19.1) | **< .001** |
| LDL Cholesterol (mg/dL) | 121.2 (38.6) | 117.0 (39.7) | 124.5 (37.7) | 0.214 |
| Triglycerides (mg/dL) | 101.5 (48.6) | 100.2 (50.5) | 102.4 (47.3) | 0.777 |
| Est TIV (cm<sup>3</sup>) | 1490 (130) | 1592 (98) | 1412 (94) | **< .001** |
| Gray Matter (% Est TIV) | 43.9 (2.1) | 42.9 (1.8) | 44.7 (1.9) | **< .001** |
| White Matter (% Est TIV) | 34.9 (2.0) | 35.1 (2.2) | 34.8 (1.9) | .362 |

**Note.** ACE III = Addenbrooke's Cognitive Examination III; Est. Adj. = Estimated cardiorespiratory fitness with Sex constant removed; METs = Metabolic Equivalents; TIV = Total Intracranial Volume. Gray and white matter volumes expressed as % of TIV. p-values from independent samples t-tests; bold indicates $p < .05$ .

Cardiovascular measures were compared against Australian norms (Nelson et al., 2024). Average brachial blood pressure indicators were at or slightly above guideline thresholds for elevated BP, with systolic BP at the 140mmHg cut-off and diastolic BP slightly below the 80mmHg threshold. Average resting heart rate was at the lower range of normal (60-100 bpm). Lipid profiles were generally within healthy ranges: HDL cholesterol was above recommended thresholds (>40 mg/dL for men, >50 mg/dL for women), LDL cholesterol was in the “optimal” range (mean 121 mg/dL, optimal <130 mg/dL), and triglycerides were in the normal range (<150 mg/dL).

Sex differences were in the expected direction (St. Pierre et al., 2022; Table 1) with males having higher diastolic BP and lower heart rate. Despite sex differences in fasting glucose and HDL cholesterol, both sex groups were in the normal range. Males also had significantly higher total intracranial volume but lower total grey matter volume (estimated as a percentage of TIV).

### Regional variability in PReFx and GM volume

Figure 1 maps PReFx values across each ROI and shows spatial variability across frontal and anterior temporo-parietal brain (see Table 2 for values). PreFx values tended to be lower across temporal regions bilaterally and left inferior frontal regions, indicative of lower arterial elasticity.

**Figure 1.**
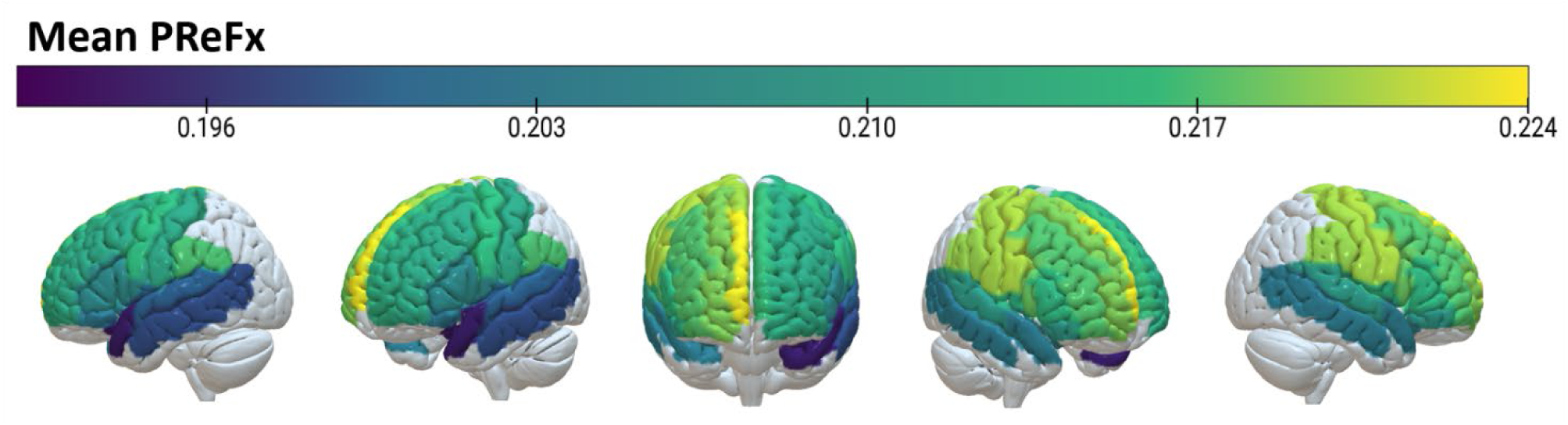
Spatial distribution of Pulse Relaxation Function (PReFx) **Note.** Mean PReFx values across 28 frontal and frontal-adjacent ROIs (N = 162). Higher values (yellow) indicate greater arterial elasticity; lower values (purple/blue) indicate reduced elasticity. Gray regions indicate areas not sampled by the optical montage. PReFx shows substantial spatial heterogeneity, with lowest values in bilateral temporal regions and left inferior frontal cortex, and highest values in right frontal and parietal cortices.

**Table 2.** Regional PReFx and gray matter volume by hemisphere.

| Region of Interest | A. PReFx |  |  |  | B. Gray Matter Volume (% EST TIV) |  |  |  |
| --- | --- | --- | --- | --- | --- | --- | --- | --- |
|  | Left Hemisphere |  | Right Hemisphere |  | Left Hemisphere |  | Right Hemisphere |  |
|  | Mean (SD) | IQR | Mean (SD) | IQR | Mean (SD) | IQR | Mean (SD) | IQR |
| **Superior Frontal Gyrus (Medial)** | 0.216 (0.044) | 0.184-0.249 | 0.223 (0.047) | 0.182-0.260 | 0.533 (0.045) | 0.500-0.566 | 0.415 (0.037) | 0.388-0.439 |
| **Superior Frontal Gyrus** | 0.213 (0.042) | 0.183-0.245 | 0.219 (0.043) | 0.181-0.251 | 0.846 (0.073) | 0.797-0.890 | 0.848 (0.073) | 0.794-0.895 |
| **Middle Frontal Gyrus** | 0.212 (0.042) | 0.180-0.249 | 0.217 (0.040) | 0.188-0.247 | 0.920 (0.745) | 0.852-0.982 | 0.921 (0.079) | 0.869-0.977 |
| **Inferior Frontal Gyrus (Opercular)** | 0.206 (0.046) | 0.172-0.241 | 0.218 (0.045) | 0.185-0.247 | 0.226 (0.021) | 0.213-0.241 | 0.278 (0.030) | 0.258-0.297 |
| **Inferior Frontal Gyrus (Triangular)** | 0.206 (0.045) | 0.170-0.243 | 0.215 (0.046) | 0.179-0.243 | 0.448 (0.047) | 0.418-0.474 | 0.382 (0.040) | 0.354-0.409 |
| **Inferior Frontal Gyrus (Orbital)** | 0.203 (0.047) | 0.173-0.236 | 0.209 (0.054) | 0.165-0.257 | 0.146 (0.015) | 0.137-0.157 | 0.146 (0.014) | 0.138-0.154 |
| **Supplementary Motor Area** | 0.217 (0.051) | 0.180-0.257 | 0.221 (0.052) | 0.177-0.262 | 0.409 (0.041) | 0.381-0.435 | 0.431 (0.037) | 0.407-0.455 |
| **Precentral Gyrus** | 0.217 (0.042) | 0.184-0.247 | 0.221 (0.038) | 0.194-0.246 | 0.663 (0.055) | 0.628-0.697 | 0.621 (0.055) | 0.576-0.663 |
| **Postcentral Gyrus** | 0.211 (0.040) | 0.181-0.240 | 0.221 (0.041) | 0.190-0.249 | 0.713 (0.061) | 0.675-0.747 | 0.663 (0.062) | 0.621-0.702 |
| **Rolandic Operculum** | 0.200 (0.041) | 0.170-0.230 | 0.214 (0.041) | 0.183-0.243 | 0.229 (0.021) | 0.216-0.242 | 0.304 (0.027) | 0.287-0.322 |
| **Supramarginal Gyrus** | 0.217 (0.045) | 0.183-0.251 | 0.220 (0.048) | 0.184-0.251 | 0.265 (0.023) | 0.249-0.280 | 0.348 (0.034) | 0.328-0.369 |
| **Middle Temporal Gyrus** | 0.198 (0.041) | 0.172-0.227 | 0.204 (0.041) | 0.173-0.228 | 1.147 (0.091) | 1.080-1.216 | 1.020 (0.071) | 0.967-1.068 |
| **Superior Temporal Gyrus** | 0.200 (0.039) | 0.172-0.225 | 0.210 (0.042) | 0.177-0.237 | 0.605 (0.053) | 0.567-0.641 | 0.632 (0.052) | 0.595-0.665 |
| **Temporal Pole (Superior)** | 0.194 (0.049) | 0.153-0.229 | 0.204 (0.050) | 0.163-0.242 | 0.349 (0.034) | 0.325-0.374 | 0.350 (0.034) | 0.325-0.371 |
| **Mean PReFx** | 0.208 (0.008) |  | 0.215 (0.006) |  |  |  |  |  |

**Note.** Values represent means (standard deviations) and interquartile ranges (IQR) for N = 162 participants. PReFx = Pulse-Relaxation Function; EST TIV = Estimated Total Intracranial Volume. Average PReFx represents mean across the 14 regions for each hemisphere.

As expected, spatially proximal regions perfused by branches of the same cerebral artery tended to be more highly correlated (e.g., left superior and middle temporal gyri, r = 0.94) compared to more distant regions (e.g., left temporal pole and right supplementary motor area, r = 0.25). Complete intercorrelation matrices are available at https://osf.io/s4vkw.

Table 2 also depicts average GM volume for each ROI.

### Associations between age, regional PReFx and regional GM volume

Figure 2 presents bivariate correlations between age, regional PReFx, and GM volume. Correlation coefficients (r) and 95% confidence intervals (10,000 iterations) for each ROI are shown in Table 3, with ROIs where confidence intervals did not cross zero highlighted in bold.

**Figure 2.**
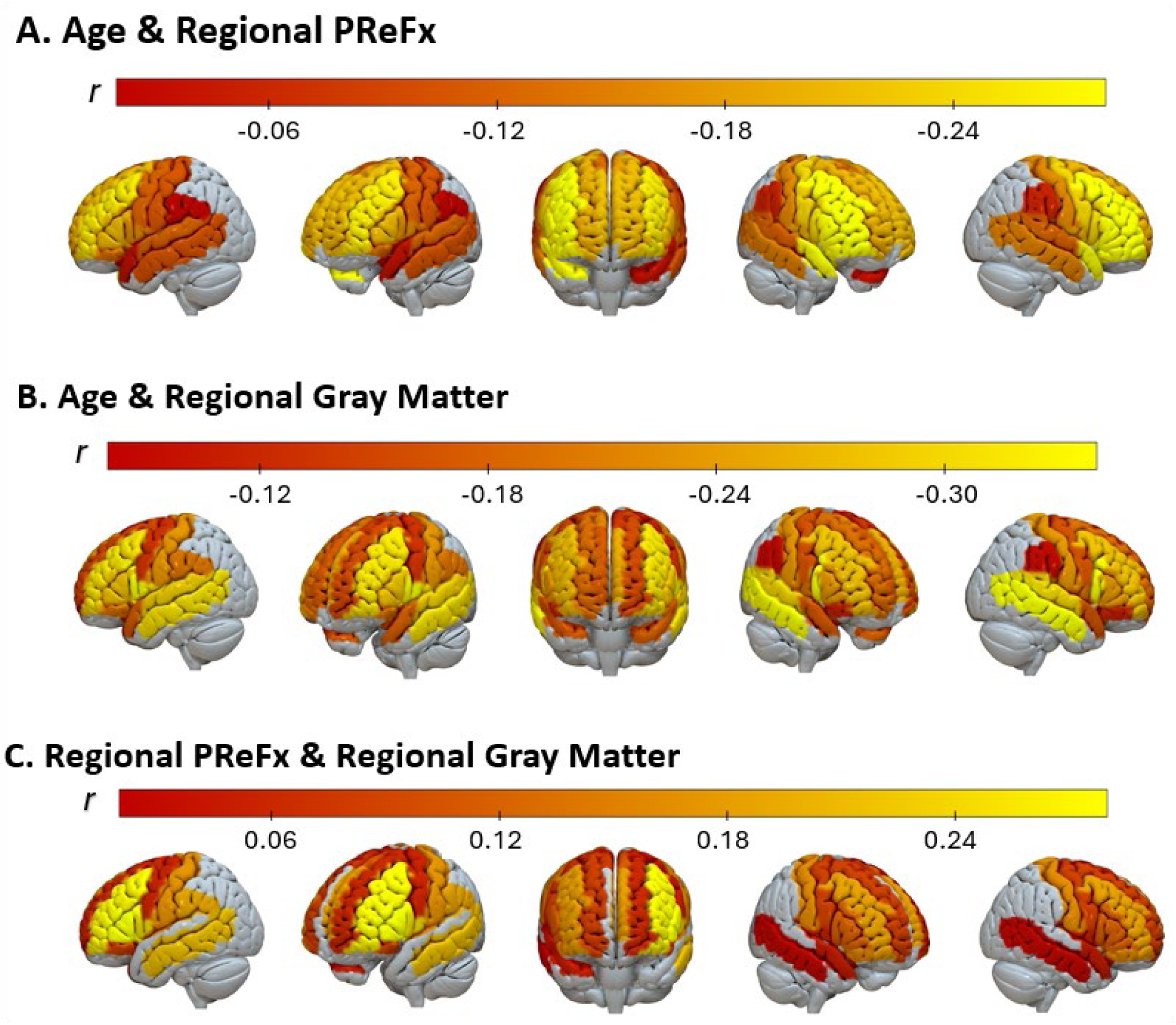
Regional correlations between age, PReFx, and grey matter volume (see Table 3 for values). **Note.** Color intensity reflects correlation magnitude: darker shades (red/orange) indicate weaker effects; brighter shades (yellow) indicate stronger effects. Panels A and B represent negative correlations (i.e., decline with increasing age). Panel C represents positive correlations (i.e., higher arterial elasticity with higher GM volume). Here five ROIs showed small negative average correlation coefficients (see Table 3) and are not depicted. Note that r-scale differs across panels.

**Table 3.** Bivariate correlations between age, cerebral arterial elasticity (PReFx), and grey matter volume by region of interest (ROI). Confidence intervals not crossing zero are bolded. ROIs highlighted represent areas used to define a mid-inferior frontal and temporal ROI for each hemisphere.

| Region of Interest | A. Age & PReFx |  | B. Age & GM Volume |  | C. PReFx & GM Volume |  |
| --- | --- | --- | --- | --- | --- | --- |
|  | Left | Right | Left | Right | Left | Right |
| **Superior Frontal Gyrus (Medial)** | **-0.19 [-0.34, -0.04]** | **-0.20 [-0.34, -0.04]** | **-0.24 [-0.38, -0.09]** | -0.16 [-0.32, 0.00] | 0.15 [-0.01, 0.30] | -0.03 [-0.18, 0.12] |
| **Superior Frontal Gyrus** | **-0.20 [-0.34, -0.04]** | **-0.18 [-0.32, -0.03]** | -0.15 [-0.31, 0.01] | **-0.18 [-0.34, -0.03]** | 0.04 [-0.11, 0.19] | 0.05 [-0.10, 0.21] |
| **Middle Frontal Gyrus** | **-0.25 [-0.39, -0.10]** | **-0.29 [-0.43, -0.13]** | **-0.29 [-0.42, -0.15]** | **-0.25 [-0.41, -0.10]** | **0.25 [0.09, 0.39]** | 0.15 [-0.01, 0.29] |
| **Inferior Frontal Gyrus (Opercular)** | **-0.15 [-0.30, -0.01]** | **-0.22 [-0.37, -0.08]** | **-0.33 [-0.46, -0.19]** | **-0.32 [-0.45, -0.19]** | **0.22 [0.07, 0.37]** | 0.10 [-0.04, 0.23] |
| **Inferior Frontal Gyrus (Triangular)** | **-0.20 [-0.34, -0.05]** | **-0.27 [-0.42, -0.11]** | **-0.26 [-0.40, -0.11]** | **-0.22 [-0.37, -0.06]** | **0.26 [0.10, 0.41]** | 0.10 [-0.05, 0.26] |
| **Inferior Frontal Gyrus (Orbital)** | -0.13 [-0.28, 0.02] | **-0.28 [-0.43, -0.12]** | **-0.19 [-0.33, -0.05]** | -0.11 [-0.27, 0.05] | 0.07 [-0.08, 0.22] | 0.14 [-0.02, 0.28] |
| **Supplementary Motor Area** | **-0.17 [-0.32, -0.01]** | -0.12 [-0.27, 0.03] | -0.13 [-0.29, 0.03] | **-0.20 [-0.36, -0.04]** | 0.09 [-0.06, 0.24] | 0.06 [-0.08, 0.20] |
| **Precentral Gyrus** | -0.11 [-0.25, 0.03] | **-0.22 [-0.36, -0.07]** | -0.12 [-0.27, 0.03] | **-0.15 [-0.30, -0.00]** | 0.03 [-0.13, 0.19] | 0.07 [-0.09, 0.22] |
| **Postcentral Gyrus** | -0.10 [-0.24, 0.03] | -0.15 [-0.31, 0.01] | **-0.21 [-0.36, -0.06]** | **-0.22 [-0.36, -0.06]** | 0.15 [-0.03, 0.33] | 0.13 [-0.02, 0.27] |
| **Rolandic Operculum** | -0.13 [-0.29, 0.02] | **-0.19 [-0.33, -0.03]** | **-0.30 [-0.43, -0.15]** | **-0.19 [-0.34, -0.02]** | 0.13 [-0.04, 0.31] | 0.03 [-0.12, 0.18] |
| **Supramarginal Gyrus** | -0.02 [-0.17, 0.12] | -0.05 [-0.21, 0.10] | **-0.19 [-0.33, -0.03]** | -0.09 [-0.24, 0.06] | **0.16 [0.00, 0.31]** | -0.07 [-0.23, 0.09] |
| **Middle Temporal Gyrus** | -0.10 [-0.25, 0.06] | -0.14 [-0.30, 0.02] | **-0.29 [-0.43, -0.14]** | **-0.34 [-0.48, -0.19]** | **0.19 [0.05, 0.34]** | 0.01 [-0.14, 0.16] |
| **Superior Temporal Gyrus** | -0.11 [-0.26, 0.05] | -0.12 [-0.28, 0.05] | **-0.27 [-0.41, -0.12]** | **-0.28 [-0.42, -0.14]** | -0.01 [-0.16, 0.15] | -0.02 [-0.16, 0.12] |
| **Temporal Pole (Superior)** | -0.03 [-0.19, 0.13] | **-0.27 [-0.41, -0.11]** | **-0.17 [-0.32, -0.02]** | **-0.16 [-0.31, -0.00]** | -0.03 [-0.17, 0.11] | 0.03 [-0.13, 0.19] |

**Note.** Pulse relaxation function (PReFx) represents cerebral arterial elasticity per region of interest; GM volume measures are proportionally adjusted gray matter volumes expressed as percentage of estimated total intracranial volume. N =162.

Figure 2A (see also Table 3A) shows that, despite the restricted 10-year age range (60-70 years), higher age was consistently associated with lower arterial elasticity, particularly in bilateral prefrontal cortex ROIs.

Increasing age was also associated with lower GM volume (Figure 2B, Table 3B), indicating greater atrophy. Correlations between age and GM volume were more widespread, evident across most ROIs bilaterally.

Correlations between regional PreFx and corresponding GM volume are shown in Figure 2C (Table 3C). Most correlations were positive, with only a few small negative correlations whose confidence intervals spanned zero. Confidence intervals excluded zero in a small number of ROIs primarily in the left-hemisphere frontal and temporal cortices (Table 3C).

### Mediation Analysis

Mediation models examined whether PReFx mediates the relationship between age and GM volume. This analysis was run at left and right frontal ROIs defined by averaging over middle frontal gyrus, and opercular and triangular parts of the inferior frontal gyrus, as well as bilateral temporal ROIs averaging over middle and superior temporal gyri and temporal pole (Figure 3).

**Figure 3.**
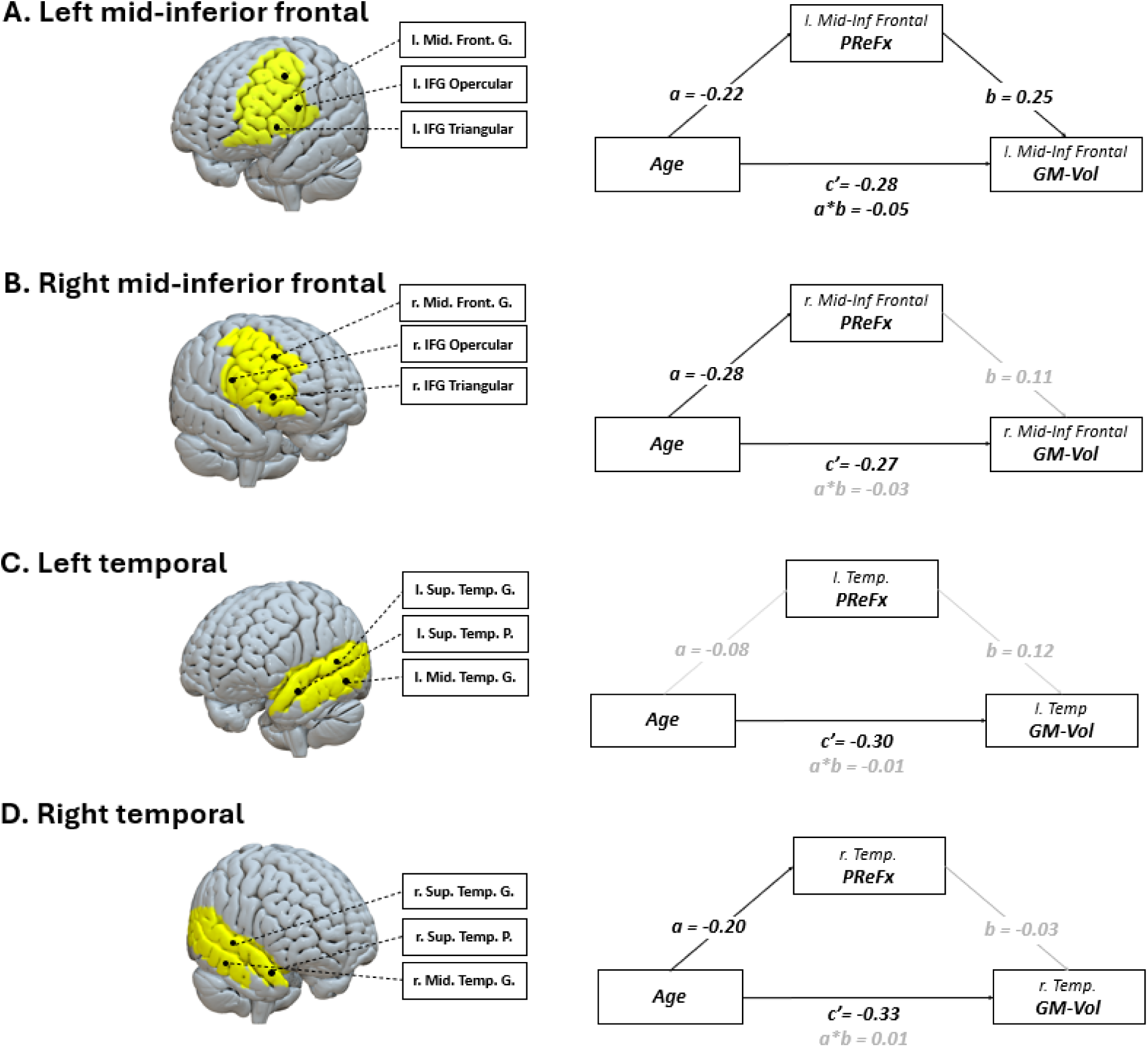
Mediation models examining prefrontal arterial elasticity (PReFx) as a mediator of age-related gray matter volume decline across four cortical regions. **Note.** Regions of interest displayed on a template brain (Rorden, 2025). Mediation path diagrams show standardised beta coefficients for each path. Significant paths and effects (*p* < .05 or 95% CI excluding zero) are shown in black; non-significant paths in gray. ROI = region of interest; l. = left; r. = right; Mid. Front. G. = middle frontal gyrus; IFG = inferior frontal gyrus; Sup. Temp. G. = superior temporal gyrus; Sup. Temp. P. = superior temporal pole; Mid. Temp. G. = middle temporal gyrus; GMVol = gray matter volume.

At the left mid-inferior frontal ROI (Figure 3A), the mediation model was statistically significant (R² = .167, F(2, 159) = 15.97, p < .001), with age and PReFx explaining 16.7% of variability in GM volume. As expected, increasing age was associated with lower PReFx (a path: β=−0.22 [−.37, −.06], p=.006), which in turn was associated with lower GM volume (b path: β=0.25 [.10, .39], p=.001). The direct effect of age on GM volume remained significant when including PReFx as a mediator (c’ path: β=−0.28 [−.42, −.13], p=.001). The indirect effect via PReFx was small but also significant (a*b path: β=−0.05 [−.11, −.01]). Therefore, PReFx in the left mid-inferior frontal ROI partially mediated the relationship between age and GM volume, with the indirect pathway accounting for approximately 16.1% of the total age effect.

In the right-hemisphere homolog (Figure 3B), the model was statistically significant overall (R² = .103, F(2, 159) = 9.08, p < .001), with age and PReFx explaining 10.3% of variability in GM volume. Again, the a-path was significant, with increasing age associated with lower PReFx (β = −0.28 [−.43, −.13], p < .001). Similarly, the direct effect of age on GM volume remained significant when adding the PReFx mediator (c’ path: β = −0.27 [−.43, −.12], p < .001). However, the b-path from PReFx to GM volume was not significant (β = 0.11 [.05, .26], p = .175), nor was the indirect effect from age to GM volume via PReFx (a*b: β = −0.03 [−.08, .01]). So, in the right mid-inferior frontal ROI, PReFx did not significantly mediate the relationship between age and GM volume.

PReFx also did not significantly mediate the relationship between age and GM volume at the two temporal ROIs. For the left temporal ROI (Figure 3C), the simple mediation model was statistically significant (R² = .112, F(2, 159) = 9.99, p < .001), with age and PReFx explaining 11.2% of variability in GM volume and a significant direct effect of age on GM volume was significant (c’ path: β = −0.30 [−.45, −.16], p < .001). However, neither the a-path (age → PReFx) nor the b-path (PReFx → GM) were significant (β = −0.08 [−.24, .07], p = .305; β = 0.12 [−.03, .26], p = .122, respectively), resulting in no indirect effect of age on GM volume (a*b: β = −0.01 [−.04, .01]).

The right temporal ROI (Figure 3D) produced a similar pattern of outcomes. The overall model was significant (R² = .109, F(2, 159) = 9.76, p < .001), as were the a-path from age to PReFx (β = −0.20 [−.35, −.05], p = .010) and the direct effect of age on GM volume (c’ path: β = −0.34 [−.49, −.18], p < .001). However, neither the b-path (PReFx → GM volume) nor the indirect effect of age on GM volume via PReFx were statistically significant (β = −0.03 [−.18, .12], p = .698; a*b: β = 0.01 [−.03, .04]).

### Moderated-Mediation Analysis

Moderated-mediation models were only fitted for the left mid-inferior frontal ROI for which PReFx partially mediated the age → GM volume relationship (Figure 4).

**Figure 4.**
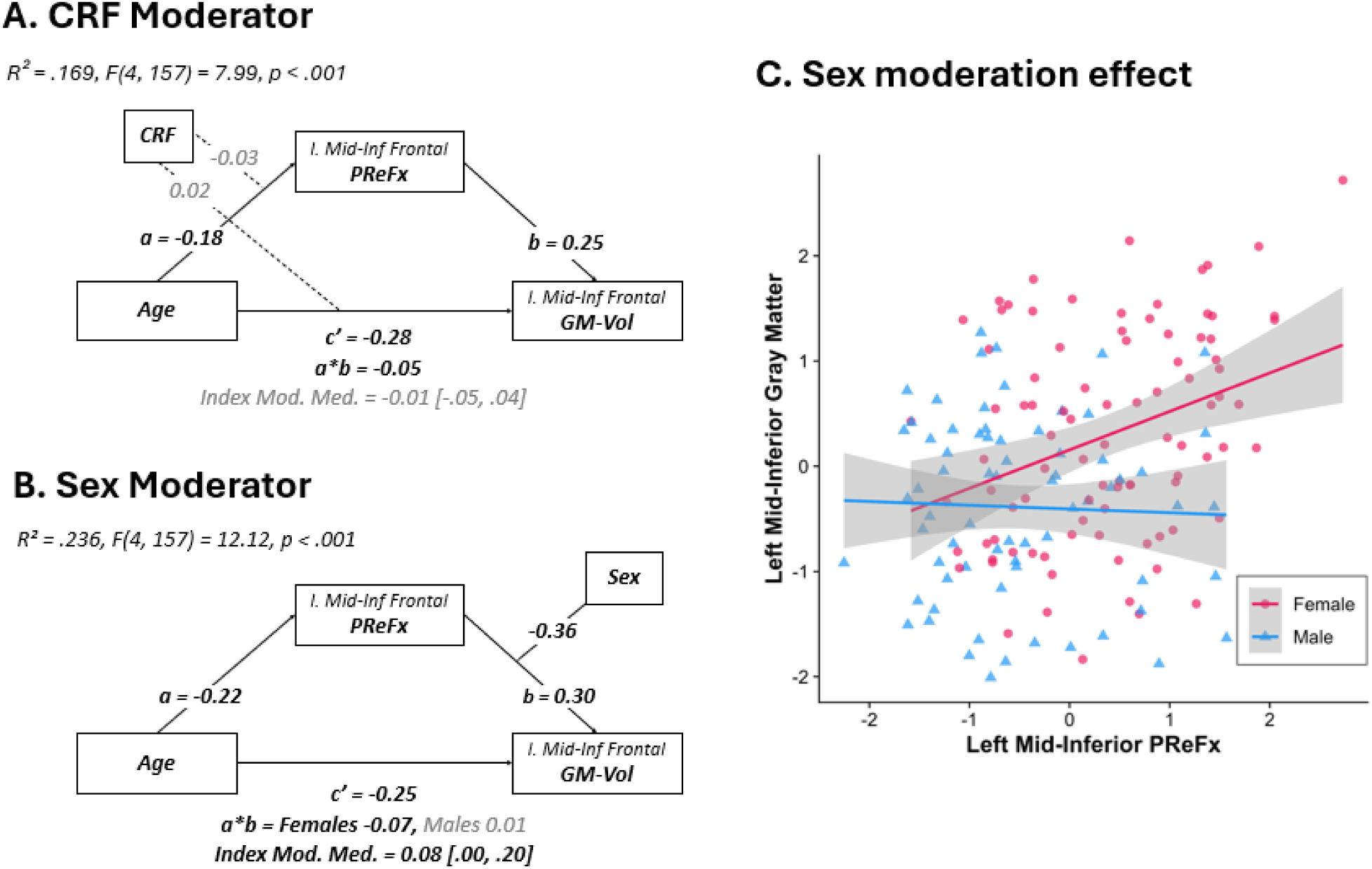
Moderated mediation models examining (A) cardiorespiratory fitness (CRF) and (B) sex as moderators of the age–PReFx–GM volume pathway in the left mid-inferior frontal region. (C) Scatter plot depicting the sex-moderated relationship between PReFx and gray matter volume (standardised values). **Note.** All path coefficients are standardised. Significant paths and effects displayed in black; nonsignificant in gray. Shaded regions represent 95% confidence intervals.

The model including CRF as a moderator of the age → PReFx (a path) and the age → GM Volume (c path) did not result in a significant index of moderated mediation (Index = −0.01, 95% CI [−.05, .04]; Figure 4A). Neither interaction term was significant (Age × CRF on PReFx: p = .740; Age × CRF on GM volume: p = .797). Thus, in this age-restricted cohort, while CRF was positively associated with PReFx (β = 0.19, p = .015), it did not moderate the relationships between age, PReFx, and GM volume.

Finally, we tested whether sex moderated the mediating effect of PReFx on the relationship between age and GM volume (Figure 4B). The index of moderated mediation was significant (Index = 0.078, 95% CI [.004, .195]), indicating that sex moderates the mediation pathway. This model explained 23.6% of variability in GM volume. Sex significantly moderated the PReFx → GM volume relationship (b-path interaction: β = -0.36, p = .024). Specifically, PReFx was positively associated with GM volume in females (β = 0.30, p = .004) but not in males (β = −0.06, p = .636). Consequently, the indirect effect via PReFx was significant for females (a*b = −0.7, 95% CI [−.15, −.01]), accounting for approximately 21% of the total age effect, but not for males (a*b = 0.01, 95% CI [−.04, .08]). As Figure 4C shows a possible outlier for females, we ran a sensitivity analysis excluding the highest PReFx value (n = 161). The indirect effect for females remained significant (a*b = −0.05, 95% CI [−0.12, −0.004]; see https://osf.io/s4vkw). Thus, PReFx partially mediated the effect of age on GM volume for females but not for males.^1^

## DISCUSSION

This is the first study to examine cross-sectional associations between regional cerebral arterial elasticity, regional grey matter volume and cardiorespiratory fitness in the frontotemporal cortex in a large age-restricted cohort of cognitively healthy and highly active older adults. Despite the narrow age range (60-70 yrs), PReFx was associated with age most strongly across frontal ROIs bilaterally, and with grey matter volume at three neighbouring ROIs over the left frontal cortex, encompassing the middle frontal gyrus and the opercular and triangular parts of the inferior frontal gyrus that are perfused by the middle cerebral artery (MCA). In a left mid-inferior frontal ROI, where age, PReFx, GM volume relationships were strongest, mediation analyses showed that arterial elasticity partially mediated the age-related decline in GM volume (∼17% of total effect), supporting the hypothesis that regional cerebral vascular changes contribute to cortical atrophy in older adults.

While cardiorespiratory fitness was associated with arterial elasticity, it did not significantly moderate the relationship between age and arterial elasticity. In contrast, sex moderated the relationship between arterial elasticity and GM volume, and this moderated mediation model more strongly explained variance in GM volume (∼24% of total effect). Specifically, higher arterial elasticity was associated with higher GM volume and mediated the relationship between age and GM volume in females, but not in males. This mediation effect was specific to the left mid-inferior frontal ROI and was not found in an adjacent ROI perfused by the same artery, or a contralateral homolog ROI perfused by the right MCA.

### Relationships between arterial elasticity and brain structure

These findings replicate, in a large cohort of cognitively healthy older adults, earlier findings by Chiarelli et al. (2017) indicating strong associations between regional arterial cerebral elasticity and GM volume, especially in the prefrontal cortex. We extend these findings by showing that, even in this age-constrained cohort, age has a significant impact on GM volume, and the relationship is partially explained by variability in arterial elasticity. Consistent with previous findings (Fabiani et al., 2014; Clements et al., 2024), higher cardiorespiratory fitness (CRF) was associated with higher arterial elasticity. However, CRF variability in this 60-70 yr age range did not significantly moderate the relationships between age and either regional arterial elasticity or regional GM volume.

Findings are also consistent with other studies using optical imaging to examine links between global and regional variation in arterial stiffness and measures of cortical structure. Mohammadi et al. (2021) reported a spatial gradient between cortical thickness and an alternative optical measure of cerebral arterial elasticity – pulse-transit-to-peak of the pulse, or the time for arterial pulse to travel from the left ventricle to the measurement point. In 30 older adults, global arterial elasticity was positively associated with global cortical thickness. These relationships were stronger bilaterally in superior, middle and inferior frontal, as well as superior and middle temporal regions, and tapered off more posteriorly. Using a similar measure of pulse transit time, Clements et al. (2024) showed that global arterial health mediated the relationship between age and white matter microstructural organization in the fornix and corpus callosum, as well as between cardiorespiratory fitness and white matter in the brainstem. These relationships were not replicated with PReFx, which measures speed of propagation of the recoil wave and therefore arterial function downstream from the point of measurement (Gratton et al., 2017). However, PReFx has been shown to mediate the relationships between age and white matter hyperintensities (Tan et al., 2019). Together, these findings extend our understanding of the link between arterial elasticity and brain structural health. They show that brain structural health is not only associated with systemic measures of arterial health, but there is also regional variability in central measures of arterial health which are associated with corresponding brain structural health.

The sensitivity of brain structural health to both global and regional variability in arterial elasticity is consistent with models proposing a hierarchical cascade of impact between vascular, brain and cognitive health with increasing age (e.g., Bateman et al., 2008; Kong et al., 2020; Stone et al., 2015; Villeneuve & Jagust, 2015). As arterial stiffening increases, pulsatile energy is transmitted more forcefully into the cerebral circulation. Sustained transmission of pulsatile force can damage the microcirculation that supplies brain parenchyma, initiating a cascade of downstream processes that are associated with blood–brain barrier disruption, impaired cerebrovascular reactivity, and regional hypoperfusion). This vascular cascade may ultimately contribute to parenchymal deterioration, manifesting as cortical atrophy and disrupted functional network organization (see Zimmerman et al., 2021 for review). The effect of arterial stiffness on cortical structure may be more severe in regions proximal to the cranial entry points of the major arteries, such as the internal carotids, where pulsatile forces are largest. Cerebral autoregulatory processes may have more time to dampen pulsatile flow at increasing distances from these entry points (Claassen et al., 2021), such that mechanical stress on the cerebral arterial wall and resultant structural changes may decrease proportionally with distance from pulsatile inflow (Mohammadi et al., 2021; see also Zarrinkoob et al., 2016 for a discussion on the pulsatility dampening factor).

The finding that PReFx partially mediated the effect of age on grey matter volume specifically at IFG and MFG is consistent with this interpretation. These areas are situated within early internal carotid artery perfusion territories and lie proximally to anterior mid-frontal watershed zones (see Kapasi et al., 2018). They have also been shown to be more vulnerable to measures of systemic arterial stiffness (Pasha et al., 2015; Celle et al., 2012). However, the left-lateralization of this effect is more challenging to interpret. We speculate that both structural and physiological asymmetries in the arterial system may have contributed to this finding. For example, studies have reported greater intima–media thickness (Luo et al., 2011) and a higher prevalence of atherosclerotic plaque (Selwaness et al., 2014) in the left carotid artery; features that may increase the pulsatile component of cerebral blood flow (see Liu et al., 2024 for review). Additionally, the right common carotid artery originates from the brachiocephalic trunk, which partially buffers pulsatile energy, whereas the left carotid artery receives direct input from the aortic arch. This anatomical asymmetry may increase the pulsatile force transmitted to the left hemisphere. Hedna et al., (2013) proposed that such vascular asymmetries might explain the elevated incidence of ischemic strokes in the left hemisphere. Notably, this difference appears largely driven by more frequent infarcts in the left middle cerebral artery (the region supplying the areas identified in the current study). However, hemispheric stroke asymmetry has been questioned, as potentially reflecting reporting bias or differences in symptom detection (see Fink, 2005 for a discussion). It is also worth noting that, although right-hemisphere associations did not reach conventional statistical thresholds in this cohort, they showed similar, albeit weaker, directional effects (Fig. 3, bottom panel).

In contrast, spatial associations were much weaker in the adjacent left temporal cortex, and this finding is inconsistent with Chiarelli et al. (2017) and Mohammadi et al. (2021), who reported significant associations between arterial and structural measures in the temporal cortex. This may arise from methodological and/or cohort differences. For instance, Mohammadi et al. (2021) report pulse-transit-to-the-peak-of-the-pulse, which like PTT reflects cumulative stiffness from the heart to the measurement site, measured cortical thickness, and excluded participants with any cardiovascular risk factors. Like Chiarelli et al. (2017), we measured PReFx which indexes regional arterial elasticity from the site of measurement to downstream peripheral resistance and GM volume. However, differences in montage coverage, sample size, age range and health status may explain differences in spatial distribution of relationships between arterial health and cortical volume. There is clearly a need for more highly powered studies to characterise the pattern of relationships and how they are affected by key cardiovascular health and other variables. However, the fact that regionally-specific associations between arterial stiffness and measures of cortical health are evident in our large cognitively healthy and highly active older ACTIVate cohort, as well as younger cohorts (e.g., Chiarelli et al., 2017), and PReFx decline precedes brain structural changes by about 10 years (Bowie et al., 2024), emphasises the potential of optical measures in early detection of changes of cerebral arterial health and prevention of later cortical and cognitive decline.

### Role of cardiorespiratory fitness

Although cardiorespiratory fitness was associated with greater arterial elasticity in the left lateral prefrontal region, level of cardiorespiratory fitness did not moderate the relationship between age and PReFx or the mediating role of PReFx in age-related grey matter volume decline. This finding was unexpected, given strong evidence linking higher level of cardiorespiratory fitness to arterial health and reduced cortical atrophy (Albin et al., 2020; Erickson et al., 2014), as well as reduced rate of age-related arterial stiffening and cortical atrophy over time (Dougherty et al., 2021; Jae et al., 2024).

This finding may, at least partly, arise from the fact that our cohort was on average highly fit, resulting in a ceiling of the moderating effect of CRF on the age to PReFx and age to GM volume relationship. Despite a mean age of 65.4 years, mean CRF was above ‘highly deconditioned’ thresholds established for middle-aged adults (Jurca et al., 2005), and their average level of moderate-to-vigorous physical activity far surpassed national guidelines (Kalamala, Ware et al., 2025; Mellow et al., 2024). Thus, our cohort seems to be comprised of *functional* to *highly functional* older adults, resulting in low sensitivity of CRF to variability in arterial and GM structural health (Montero et al., 2017). Nevertheless, our cohort did show substantial variability in presence of cardiovascular risk factors, and this was associated with differences in PReFx (see Johnson et al., in preparation). Further longitudinal work with larger samples is needed to characterise the interactive relationships between these measures of cardiovascular health.

### Role of biological sex

Biological sex moderated the arterial elasticity-cortical volume pathway, with PReFx mediating age-related grey matter volume decline in females but not males. This pattern may reflect sex differences in the timing and trajectory of vascular aging. Bowie et al. (2024) showed that global cerebral arterial stiffness accelerates sharply in women around age 50 and precedes structural brain changes by approximately a decade. Our 60–70-year cohort may therefore capture a period where earlier arterial stiffening is now manifesting as cortical decline in females. This interpretation aligns with broader reports of stronger vascular-structural relationships in females (Armstrong et al., 2019; Reas et al., 2021) and accelerated systemic arterial stiffening following menopause. However, we did not collect data regarding menopausal status or hormone replacement therapy, limiting our ability to test these mechanisms directly. Further research with comprehensive reproductive health measures is needed to clarify the pathways linking female-specific arterial stiffening to structural brain changes.

### Conclusions

Regional cerebral arterial elasticity partially mediated age-related decline in left prefrontal grey matter volume in cognitively healthy older adults. This mediation was regionally specific and sex-dependent, emerging only in females. Cardiorespiratory fitness was associated with higher arterial elasticity but did not moderate the relationship between age, vascular and brain structural health in this active cohort. These findings support vascular cascade models and suggest arterial stiffening contributes to cortical atrophy. The sex-specific pathway indicates that females may be particularly vulnerable to vascular-mediated brain changes in older adulthood. Regional cerebral arterial elasticity measures may enable early detection of vascular changes preceding structural decline.

## Author Contributions

NW: Conceptualization, Formal analysis, Writing – Original Draft, Writing – Review and Editing, Investigation; JJ: Formal analysis, Writing – Review and Editing; MF: Writing – Review and Editing, Supervision; PM: Writing – Review and Editing, Supervision; KL: Writing – Review and Editing; CB: Formal analysis, Review and Editing; DB: Formal analysis; GG: Writing – Review and Editing, Supervision; AS: Conceptualization, Writing – Review and Editing, Funding acquisition; FK: Conceptualization, Formal analysis, Writing – Review and Editing, Funding acquisition, Supervision. All authors contributed to the article and approved the submitted version.

## Acknowledgements

We thank Michael Breakspear for consulting on brain imaging protocols and analyses, and Felicity Simpson, Nathan Tran, Montana Hunter, Alexandra Wade, Maddison Mellow, Louise Massie, Kate Dyer, Karen Wilson, Gemma Mieko Furuhashi, Helen Nicholas, Riley Jackson, Teigan Cotterill, Mahmoud Abdolhoseini, and Fayeem Aziz for valued contribution towards data collection and study coordination. We also thank the HMRI Research Volunteer Register for partial recruitment of participants at the Newcastle site.

## Data Availability

Due to the ongoing longitudinal nature of the ACTIVate study, the dataset analyzed during this study is not publicly available. However, it may be obtained from the corresponding author upon reasonable request. The supplementary results are available at the OSF repository: [https://osf.io/s4vkw].

## Footnotes

1 We also tested sex moderation of both b and c’ paths (PROCESS Model 15). The index of moderated mediation was marginally significant, and the model accounted for roughly the same variance as Model 3. Sex did not moderate the direct effect (a*b). Model 15 can be viewed at https://osf.io/s4vkw.

